# Infectivity of a pathogenicity-attenuated *Chlamydia muridarum* mutant in the genital tract

**DOI:** 10.1101/2024.12.19.629466

**Authors:** Caiting Li, Zhaoyang Liu, Yaoqin Hua, Chunguang Ma, Guangming Zhong

**Affiliations:** Department of Dermatology, The First Affiliated Hospital, Sun Yat-sen University, 58 Zhongshan 2^nd^ Road, Guangzhou, Guangdong 510080, China; Department of Microbiology, Immunology, and Molecular Genetics, University of Texas Health Science Center at San Antonio, 7703 Floyd Curl Drive, San Antonio, Texas 78229, USA

**Keywords:** IntrOv, Oral vaccine, Infectivity, Pathogenicity, Genital tract

## Abstract

A *C. muridarum* mutant designated as intrOv was evaluated as an *intr*acellular *O*ral vaccine *v*ector because it can induce protection in the genital tract following oral inoculation but does not elicit genital pathology following intravaginal infection. However, the mechanism of intrOv’s attenuation is unclear. Here we report that few live organisms were recovered from vaginal swabs during the early stage of intrOv intravaginal infection in mice. At a low inoculating dose, an isogenic wild-type control strain established a productive infection, while intrOv failed to do so. Although a higher inoculating dose allowed intrOv and its control to productively infect mice, fewer live intrOv than the control organisms were recovered from the lower genital tract tissues on day 3 post-infection. By day 7, animals infected with intrOv or the control shed similar numbers of live organisms, suggesting the intrOv’s deficiency on day 3 was transient. Consistently, intrOv reduced invasion of epithelial cells but maintained as robust intracellular replication as its control. Our results correlate intrOv’s delay in infecting the lower genital tissues and reduction in invading epithelial cells with its attenuation in genital pathogenicity, laying the foundation for further revealing the mechanisms of the intrOv’s attenuation in pathogenicity during genital tract infection.

## Introduction

*Chlamydia trachomatis* is the major cause of sexually transmitted bacterial infections, potentially leading to pathological sequelae in the upper genital tract (1–4). The precise mechanism of chlamydial pathogenicity in the upper genital tract remains unclear. The mouse-adapted chlamydial species, *C. muridarum*, can induce pathology in the mouse genital tract similar to that induced by *C. trachomatis* in women and is thus used to investigate chlamydial pathogenicity (5). All chlamydial organisms share a biphasic life cycle that alternates between the small (∼200 nm) extracellular infectious elementary body (EB) and larger (∼600 nm) intracellular replicating reticulate body (RB) morphologies. The natural target cells of chlamydial infection are mainly epithelial cells. The first steps of chlamydial infection of an epithelial cell involve an EB attaching and entering a host cell via various mechanisms, including glycosaminoglycan binding [3–6], type III secretion system (T3SS)-associated adherence [7], clathrin-mediated endocytosis [8,9], and other attachment/entry mechanisms to be discovered. Chlamydial attachment and entry can be enhanced *in vitro* by pretreating host cells with diethylaminoethyl (DEAE)-dextran (6) and/or centrifugation (7). Following entry, an EB transitions to an RB for intracellular replication, which simultaneously transforms the chlamydial organism-laden endocytic vesicle into a parasitophorous vacuole, called an inclusion. After multiple rounds of intracellular replication via binary fission, progeny RBs transform back into EBs to exit the infected host cell and infect a new host cell. Chlamydial intracellular biosynthesis must compete with the host cells since inhibition of host cell protein synthesis using cycloheximide has been shown to promote chlamydial growth (6). The progeny chlamydial organisms can exit the infected cells via multiple mechanisms, including cell/inclusion lysis and inclusion extrusion (8). Chlamydial infection may directly damage the epithelial cells and trigger host inflammatory responses, which are considered critical virulence determinants in chlamydial pathogenicity (9).

Intravaginal inoculation with *Chlamydia muridarum* can induce hydrosalpinx, a hallmark of tubal infertility detected in women with chlamydia (3, 4), in many different strains of mice (10). By taking advantage of gene knockout, the mouse genital tract infection model has identified several host factors that can regulate chlamydial pathogenicity in the genital tract, including lymphocytes, cytokines, and other molecules. For example, antigen-specific CD8^+^ T cells (11–13) promote while non-specific CD8^+^ T cells (14, 15) reduce chlamydial induction of hydrosalpinx. TNFα receptor signaling, but not TLR2 signaling, may contribute to chlamydial pathogenicity in the genital tract (16, 17). Additionally, the complement component C5 but not C3 was found to be essential for mice to develop hydrosalpinx in response to *C. muridarum* infection (18).

The *C. muridarum* infection model has also been used to identify chlamydial virulence factors contributing to the development of upper genital tract sequelae (19). For example, the plasmid-encoded Pgp3 and Pgp5 are important for *C. muridarum* to induce upper genital tract disease (20, 21). Deficiency in Pgp3 or Pgp5 reduces ascending infection and chronic inflammatory infiltration in the upper genital tract. *C. muridarum* mutants with mutations in chromosomal genes are attenuated in genital pathogenicity (22–24). Some mutants were identified by screening for genetic variants induced by chemical mutagenesis (25), while other spontaneous mutants were identified by serial *in vitro* passage (23). Some of these mutants, besides their lack of genital pathogenicity following an intravaginal inoculation, can also induce protective immunity in the genital tract following an oral inoculation (24, 26). One of these *C. muridarum* mutants was designated as intrOv or intracellular oral vaccine vector. IntrOv carries a substitution mutation in *tc0237*, that replaces a Q residue at the 117 position with E (TC0237Q117E), and introduces a nonsense mutation in *tc0668*, resulting in a premature stop codon that truncates the 408aa protein of TC0668 at G216 (TC0668 G216*). Unlike its control, intravaginal inoculation of mice with 2 x 10^5^ IFUs of intrOv failed to induce hydrosalpinx. Surprisingly, mice infected with intrOv or the control shed similar numbers of infectious organisms (22). Thus, the mechanism of the intrOv’s attenuation in pathogenicity remains unclear. Given the critical roles of chlamydial infectivity and ascending infection in chlamydial pathogenicity, it is necessary to reevaluate the intrOV’s infectivity in the genital tract tissues.

In this study, we compared the infectivity of intrOv and its isogenic wildtype control in the mouse genital tract by monitoring the live chlamydial organism yields recovered from both vaginal swabs and six different genital tissue segments and at different times after the intravaginal infection with different inoculum doses. These experiments revealed a reduced infectivity of intrOv in the genital tract. The peak level of shedding live intrOv organisms detected in vaginal swabs was delayed compared to the control, following intravaginal inoculation with 2 x 10^5^ IFUs. Live intrOv yields recovered in swabs on days 1 and 3 were significantly lower than the control. However, by day 7, the intrOv titer reached the level of its control. At a lower inoculum, the infection course of intrOv was significantly reduced and shortened. At the lowest inoculum tested, intrOv failed to establish productive infection, whereas the control remained infectious. IntrOv also displayed a reduced invasion of epithelial cells *in vitro.* These observations have provided direct evidence that intrOv is attenuated in infecting the female genital tract, which may contribute to its attenuated genital pathogenicity.

## Results

### 1. IntrOv is less efficient at establishing an infection course in the female genital tract

We previously demonstrated that intrOv failed to induce hydrosalpinx (22, 23). However, the infections caused by intrOv and its control were similar when we used an inoculum of 2 x 10^5^ inclusion forming units (IFUs). To investigate the mechanism of intrOv attenuation, we compared the infection time courses of intrOv and the control using varying inoculating doses (Fig. 1). Infections with the control resulted in infections that lasted 3 to 4 weeks regardless of inoculum doses tested. The infection peaks were postponed and moved to the right as the initial inoculum dose decreased. Specifically, live organism shedding peaked on day 3, day 7, or days 7 to 14 when inoculums of 2 x 10^5^ IFUs, 2 x 10^4^ IFUs, and 2 x 10^3^ IFUs were used, respectively. In contrast, shedding of live intrOv peaked on day 7 when we used an inoculum of 2 x 10^5^ IFUs, similar to what we observed following inoculation with 2 x 10^4^ IFUs of the control organisms, showing the ability of intrOv to establish infection is reduced by at least 10-fold. This defect was more pronounced at lower inocula. At 2 x 10^4^ IFUs, shedding of intrOv was significantly reduced, and the overall infection time course was shorter. At 2 x 10^3^ IFUs, most mice inoculated with intrOv did not become infected, whereas all mice inoculated with the control became productively infected.

**Fig. 1.**
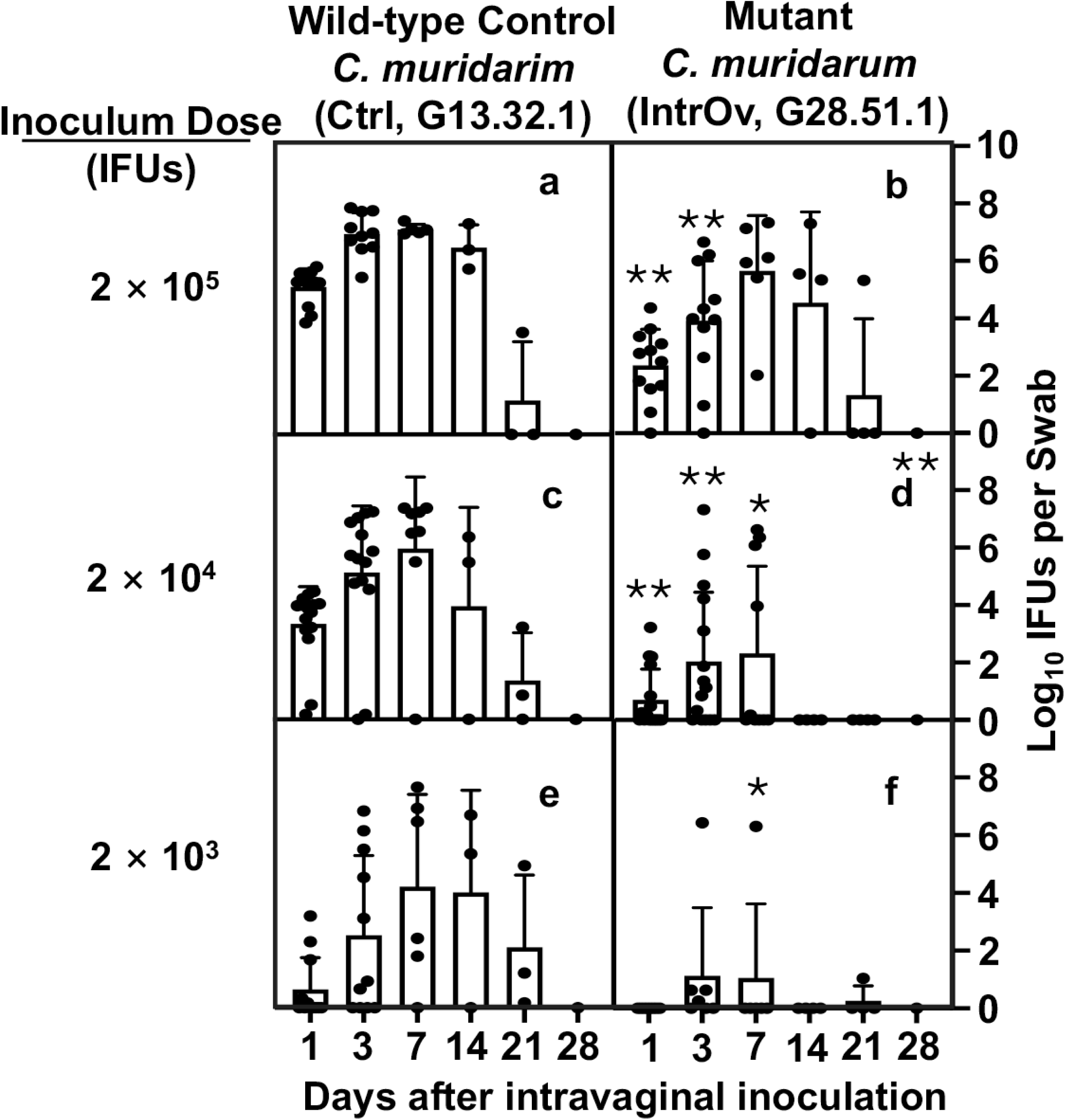
Time courses of live chlamydial organism shedding in vaginal swabs following intravaginal inoculation with wild-type control or mutant *C. muridarum*. Groups of C57BL/6J mice (n=4 to 12) were intravaginally inoculated with wild-type control (Ctrl, clone G13.32.1, panels a, c, & e) or mutant (intrOv, clone G28.51.1, b, d & f) *C. muridarum* at the dose of 2 x 10^5^ (a & b), 2 x 10^4^ (c & d), or 2 x 10^3^ (e & f) inclusion forming units (IFUs) as listed on the left, and monitored for live chlamydial organism shedding in vaginal swabs on days 1, 3, 7, and weekly after that as indicated along the X-axis. The results were expressed as Log_10_ IFUs per swab shown along the Y-axis. The data were compiled from 3 or more independent experiments. The live organism shedding time courses were converted into Area-Under-Curves (AUCs) for comparison between the Ctrl and intrOv groups using ANOVA (all groups) and Wilcoxon (each dose), respectively. Log_10_ IFUs detected from different time points under a given inoculating dose were similarly compared using ANOVA (all time points under a given dose) and Wilcoxon (each time point). *p<0.05, **p<0.01, 2-tailed Wilcoxon.

Reduced shedding of live organisms from intrOv-infected animals was most evident in the first-week post-inoculation. Fewer intrOv compared to control organisms were harvested on days 1 and 3. However, the number of live organisms recovered by day 7 did not differ between the intrOv and the control when we used an inoculum of 2 x 10^5^ IFUs, the same dose employed in our prior studies (22, 23).

### 2. IntrOv is significantly delayed in establishing productive infection in the lower genital tract tissues

To identify the tissues where intrOv is deficient in establishing infection, the tissue distribution of live intrOv and control organisms was compared. Since intrOv shedding was reduced 3 days post-infection, we focused on this time point (Fig. 2). Regardless of the infection conditions, the highest numbers of live chlamydial organisms were detected in lower genital tract (LGT) tissues, including the lower vagina (LV), upper vagina (UV) and ectocervix (EC). LGT was the inoculating site, and more IFUs were recovered from the LGT tissues than were inoculated, indicating that the LGT tissues were productively infected 3 days after inoculation. Live organisms were detected in the UGT segments of most mice inoculated with the control, suggesting that the control was able to ascend from the LGT to the UGT within 3 days of inoculation. Overall, recovery of live intrOv was reduced compared to the control in most genital tissues, and the differences were most significant with lower inocula.

**Fig. 2.**
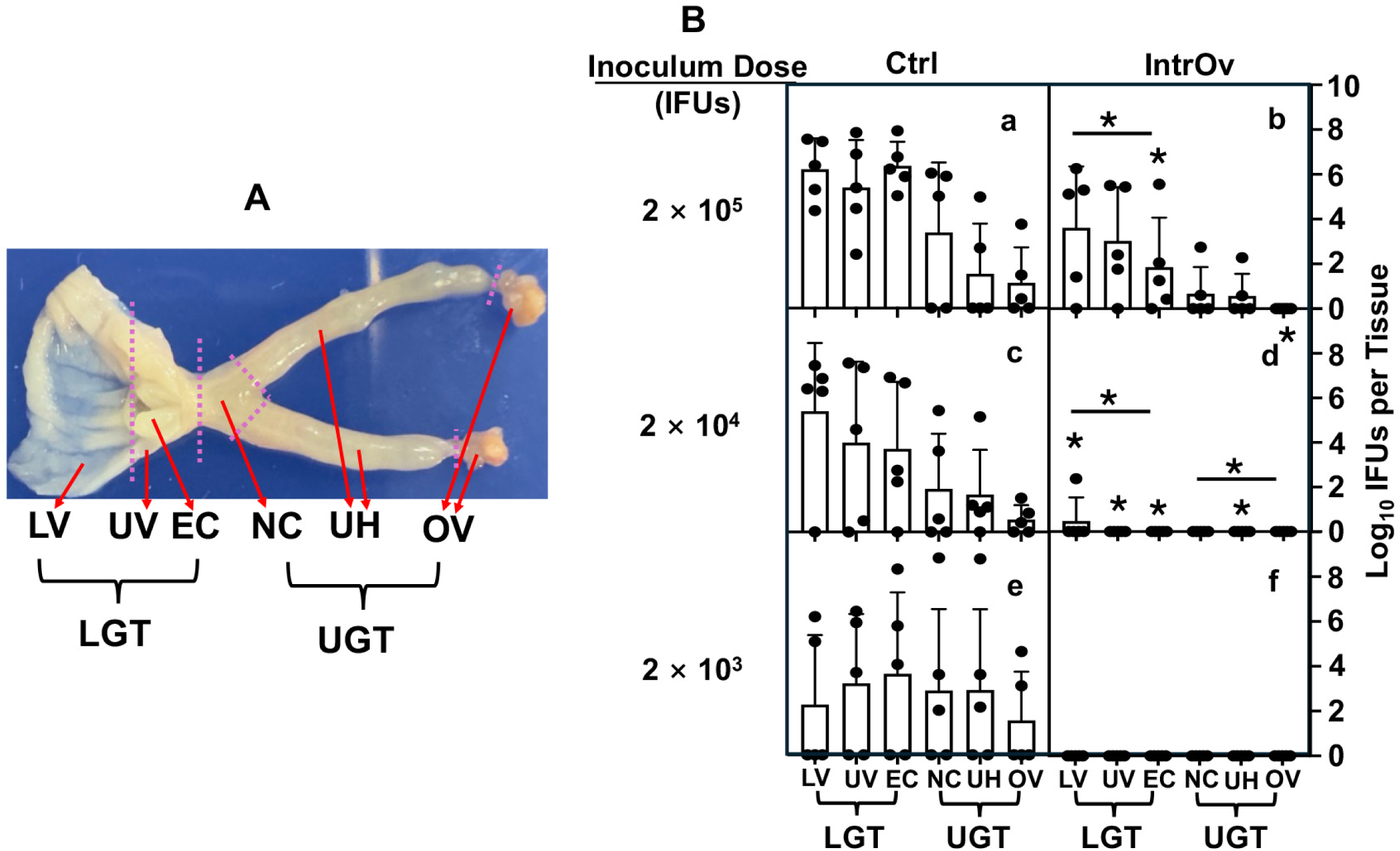
Distribution of live chlamydial organisms in genital tissues harvested 3 days following intravaginal infection. (A) A female mouse genital tract is dissected into six segments from the lower genital tract (LGT) to the upper genital tract (UGT) tissues. The LGT includes the lower vagina (LV), upper vagina (UV), and ectocervix (EC) while the UGT includes the endocervix (NC), uterine horns from both sides (UH), and Oviducts/ovaries from both sides (OV). Note that the UV segment tissue was trimmed off around the EC tissue as indicated. (B) Groups of C57BL/6J (n=5) were intravaginally inoculated with wild-type control (Ctrl, clone G13.32.1, panels a, c, & e) or mutant (intrOv, clone G28.51.1, b, d & f) *C. muridarum* at different inoculum doses as indicated on the left. On day 3 after infection, mice were sacrificed for monitoring live chlamydial organisms in 6 different genital tract tissues as listed along the X-axis. The results were expressed as Log_10_ IFUs per tissue shown along the Y-axis. The data were compiled from two independent experiments. The Log_10_ IFUs recovered from each genital tract tissue were compared between the Ctrl and intrOv groups using ANOVA (all tissues) and Wilcoxon (each segment or tissue), respectively. The number of live intrOv recovered from different genital tissues was lower than that of the Ctrl, with the most significant differences detected in the LGT and under lower inoculation doses. The lack of statistical difference in the recovered IFUs between Ctrl and intrOv at 2 x 10^3^ (e vs. f) was largely due to the high intragroup variation and the limited number of live organisms recovered from each tissue. *p<0.05, **p<0.01, 2-tailed Wilcoxon.

By day 7 post-inoculation, live organism recoveries were more similar in intrOv- and control chlamydial organism-infected animals. Even when the inoculum was reduced to 2 x 10^4^ IFUs, significant numbers of intrOv organisms were recovered from all of the genital tissues at 7 days, although the intrOv organism numbers were significantly lower than in animals inoculated with the control.

### 3. Ascension of intrOv from the lower to the upper genital tract is reduced compared to the control

We developed an index to compare the ascending efficiency of intrOv and the control (Fig. 4). The index was calculated using the number of live organisms recovered from a specific LGT segment or all 3 LGT segments as the denominator to divide the number of live organisms recovered from the UGT. In mice infected with the control, the ascending indices were below 0.5 in most mice on day 3 post-infection but closer to 1 in most mice by day 7. Assuming the LGT and UGT are equally susceptible to chlamydial infection, these observations suggest that the control had begun to ascend by day 3 post-infection, and the ascending was mostly complete by day 7. The ascension index was significantly reduced in mice infected with 2 x 10^5^ or 2 x 10^4^ IFUs of intrOv compared to the control, suggesting intrOv is not only less efficient in infecting the lower genital tract tissues, but is also less efficient in ascending across the cervical barrier. By day 7, the ascension of intrOv and the control only differed when we used low inoculum. Overall, these observations demonstrated an impaired ascension of intrOV, but this defect could be overcome at higher infectious inocula.

**Fig. 3.**
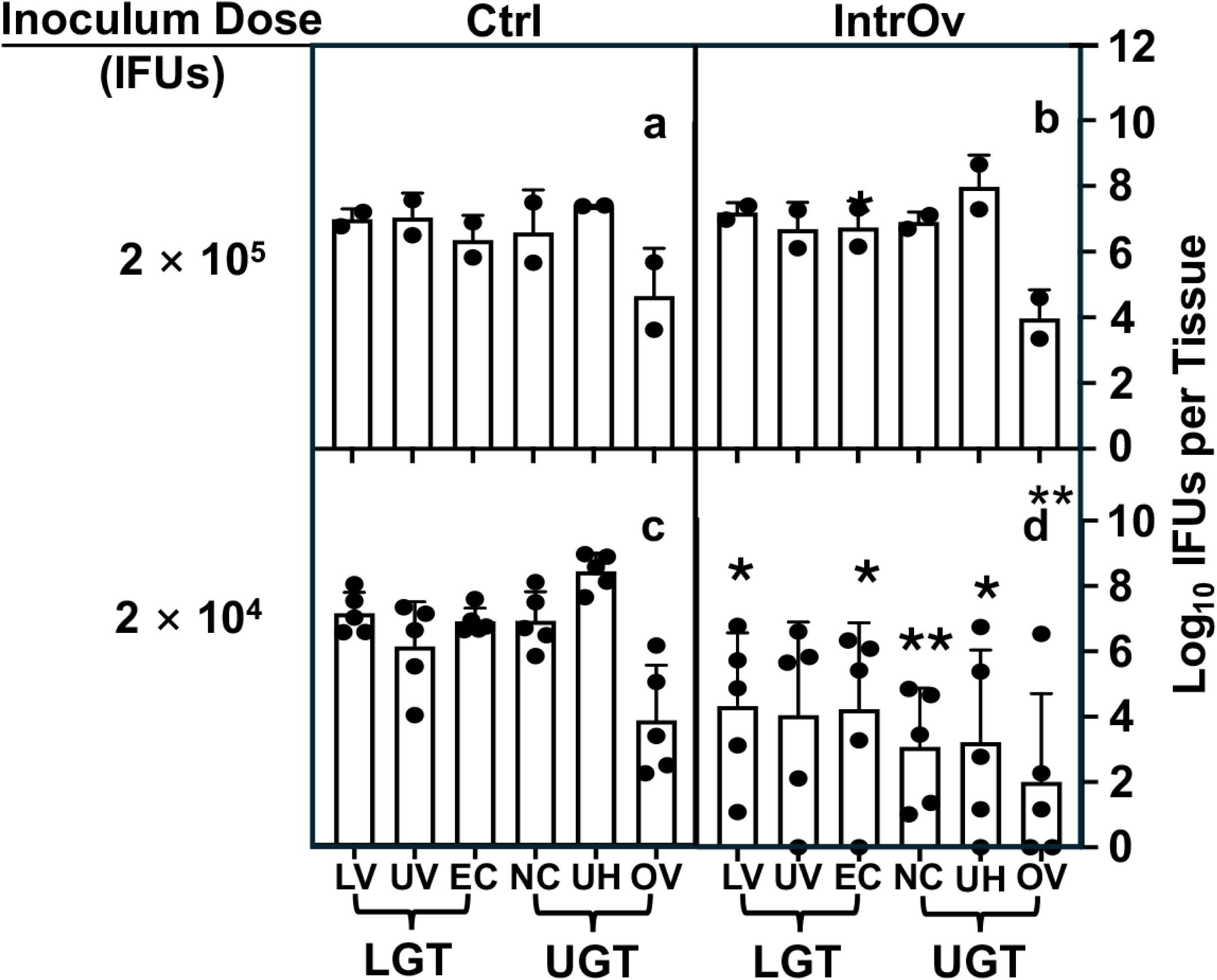
Distribution of live chlamydial organisms in the genital tract tissue harvested on day 7 following intravaginal inoculation. Groups of C57BL/6J were intravaginally inoculated with wild-type control (Ctrl, clone G13.32.1, panels a & c) or mutant (intrOv, clone G28.51.1, b & d) *C. muridarum*, and sacrificed for titrating live chlamydial organisms in six different genital tract tissues as described in Fig. 2 legend except that mice were sacrificed on day 7 after the intravaginal inoculation (n=5 for the 2 x 10^4^ inoculum dose groups, c & d and n=2 for 2 x 10^5^ groups, a & b). The results were expressed as Log_10_ IFUs per tissue shown along the Y-axis. The data were compiled from two independent experiments. The Log_10_ IFUs recovered from each genital tract tissue were compared between Ctrl and intrOv (the 2 x 10^4^ groups only) using ANOVA (all tissue segments) and Wilcoxon (each segment), respectively. *p<0.05, **p<0.01, 2- tailed Wilcoxon.

**Fig. 4.**
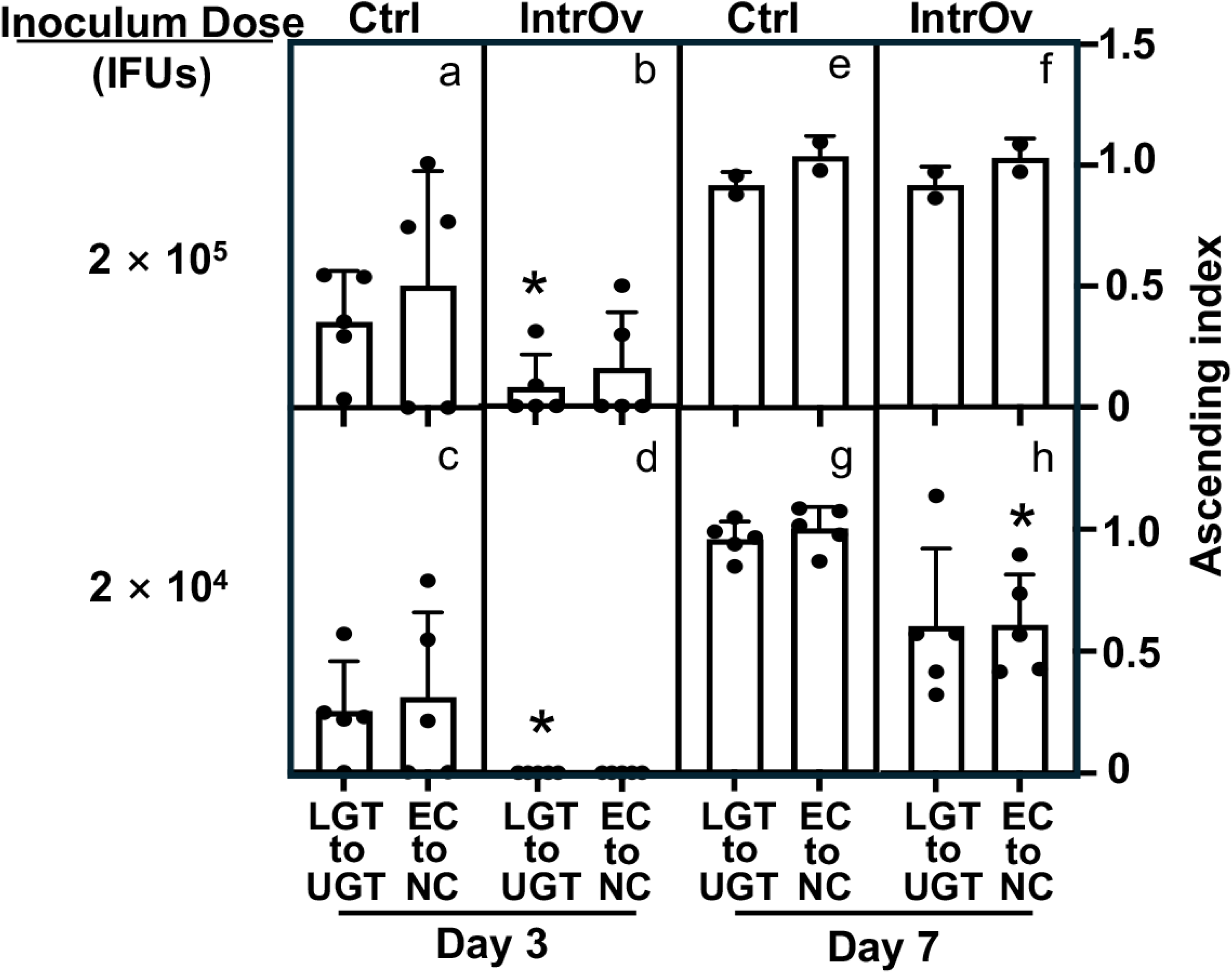
Comparison of ascending index between the Ctrl and intrOv groups. The ascending index was calculated using log_10_IFUs recovered from different genital tissues harvested on days 3 (panels a-d) and 7 (e-h), as described in Fig. 2 & 3 legends. To calculate the index of chlamydial ascending from the ectocervix to the endocervix (EC to NC, right column in each panel) of a given mouse, the Log_10_ IFUs recovered from the mouse EC was used (as the denominator) to divide the Log_10_ IFU recovered from the NC of the same mouse. To calculate the ascending index for chlamydial spreading from the lower genital tract to the upper genital tract (LGT to UGT, left column in each panel), the total Log10 IFUs from all three LGT segments of a given mouse were used to divide those from all three UGT segments of the same mouse. The index (mean and standard error shown along the Y-axis) was compared between the Ctrl and intrOv groups inoculated with a dose of 2 x 10^5^ (a, b, e, f) or 2 x 10^4^ (c, d, g, h) and measured on days 3 (a-d) or 7 (e-h) using ANOVA and Wilcoxon, respectively. *p<0.05, 2-tailed Wilcoxon.

### 4. Attachment/Entry of IntrOv is delayed, but upon entry, it develops with normal kinetics

To investigate why it takes longer for intrOv to establish productive infection of the LGT and ascend from the LGT to the UGT, we evaluated intrOv infection of cervical epithelial cells using HeLa cells as the model. DEAE-dextran treatment and centrifugation have been routinely used to enhance chlamydial attachment to and entry of host cells (6, 7), we compared the dependence of intrOv versus its control on the DEAE-dextran and centrifugation (Fig. 5). We did not observe any difference in their dependence on DEAE-dextran treatment, but a big difference in their dependence on centrifugation was found. Specifically, the % of dependence on DEAE-dextran was <3% for the control and <2% for intrOv, while the % of dependence on centrifugation was ∼30% for the control and >60% for intrOv (p<0.01). Although centrifugation significantly elevated the infectivity of both the control and intrOv, intrOv was more dependent on centrifugation for invading HeLa cells. The intrOv’s dependence on centrifugation was maintained when MOI was reduced to 0.1. These observations suggest that intrOv is less efficient in invading (attaching and entering) HeLa cells than the control.

**Fig. 5.**
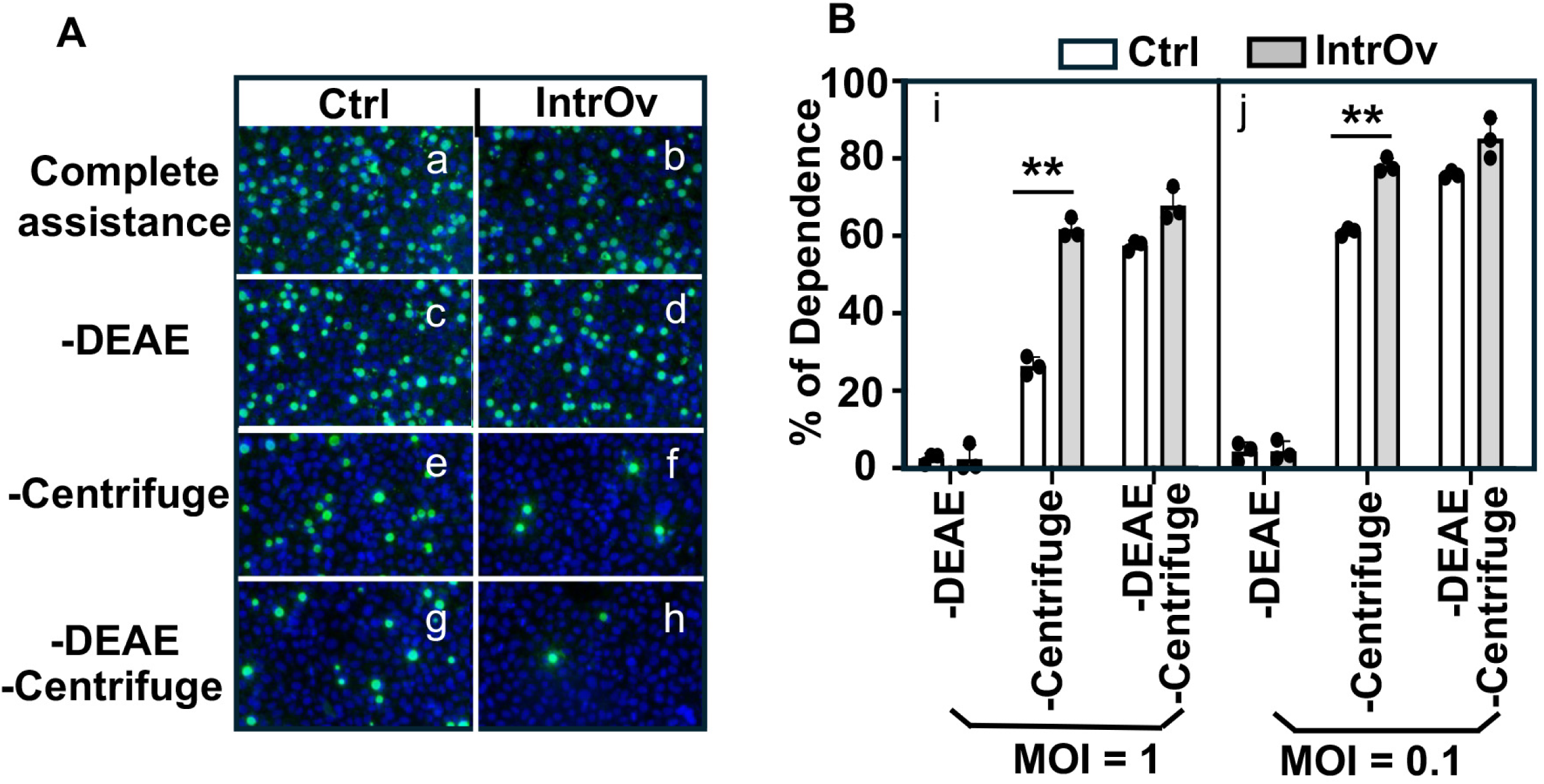
Comparison of dependence on the assisted attachment and entry between Ctrl and intrOv. (A) HeLa monolayers were inoculated with the Ctrl (panels a, c, e, & g) or intrOv(b, d, f & h) *C. muridarum* with both DEAE pretreatment of HeLa monolayers and centrifugation to assist the attachment and entry (complete assistance, a & b), no DEAE (c & d), no centrifugation (e & f) or neither DEAE nor centrifugation (g & h). After washing to remove extra inoculum, the HeLa monolayers were replenished with a warmed medium containing cycloheximide and incubated overnight. The infected monolayers were processed for immunofluorescence labeling of chlamydial inclusions (green) and host cell nuclei (blue). Representative images acquired from an experiment with an infection dose at MOI=1 were shown. As shown in B, the number of chlamydial inclusions was counted from each well and used to calculate the % of dependence on a given assisted condition. The experiments were performed under an MOI of 1 (i) or 0.1 (j). Data came from 3 independent experiments. The % of dependence was compared between the Ctrl and intrOv groups using ANOVA and Wilcoxon, respectively. *p<0.05, **p<0.01, 2-tailed Wilcoxon.

Since cycloheximide, a eukaryotic protein synthesis inhibitor, has been used to promote chlamydial intracellular growth without affecting chlamydial attachment or entry (6), we next compared the dependence of intrOv versus its control on cycloheximide-enhanced intracellular growth (Fig. 6). Surprisingly, there was no significant difference in the % of dependence between intrOv and its control regardless of the attachment/entry conditions or the MOIs used.

**Fig. 6.**
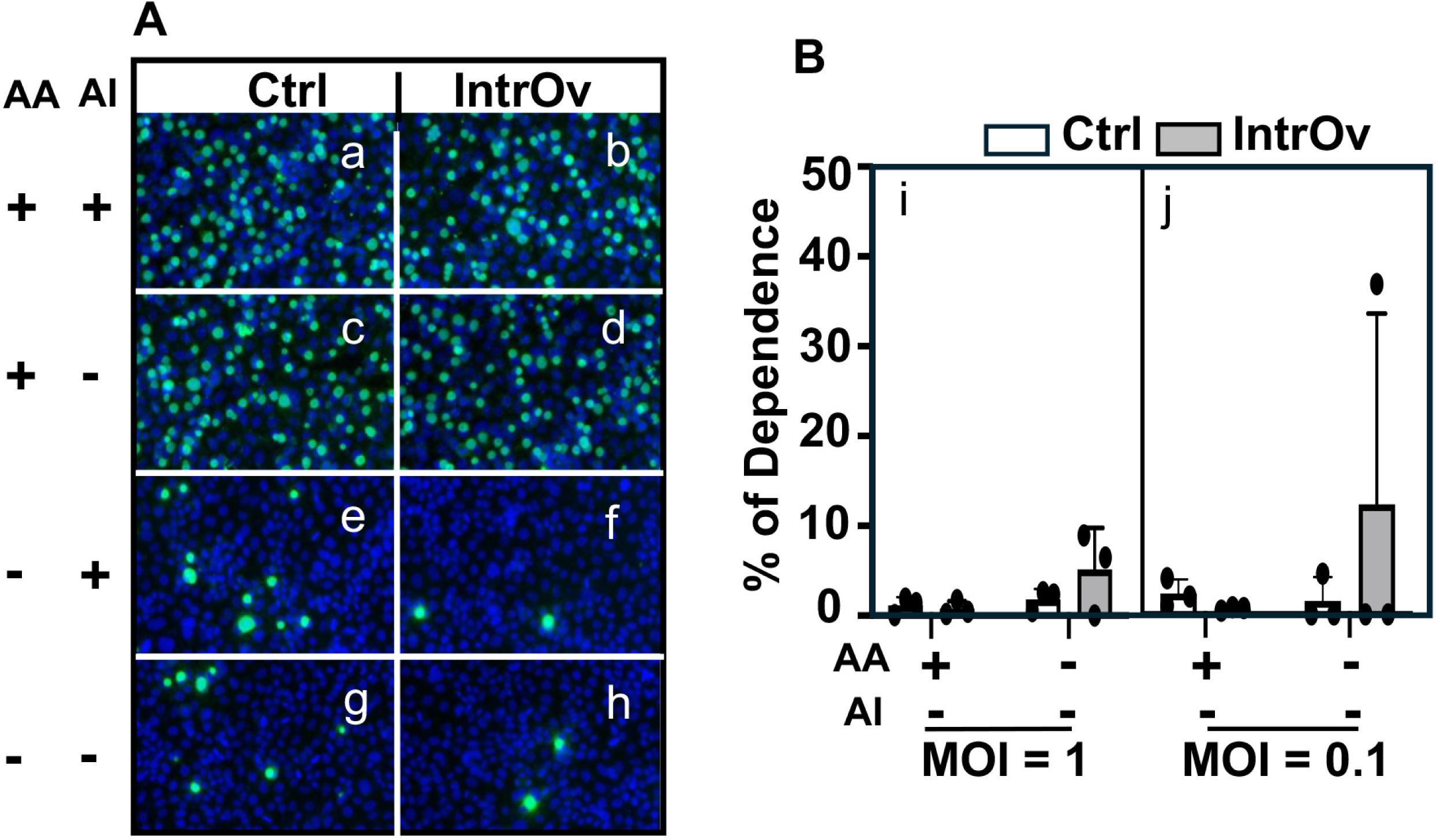
Comparison of dependence on the assisted intracellular growth between Ctrl and intrOv. (A) HeLa monolayers were inoculated with wild-type control (Ctrl, clone G13.32.1, panels a, c, e, & g) or mutant (intrOv, clone G28.51.1, b, d, f & h) *C. muridarum* with (a-d) or without (e-h) the DEAE-dextran and centrifugation to assist attachment and entry (AA). The infected cultures were incubated overnight in a medium with (a, b, e, & f) or without (c, d, g, & h) cycloheximide to assist intracellular growth (AI). The infected monolayers were processed for immunofluorescence labeling of chlamydial inclusions (green) and host cell nuclei (blue). Representative images acquired from an experiment with an infection dose at MOI=1 were shown. As shown in B, the number of chlamydial inclusions was counted from each well and used to calculate the % of dependence on intracellular growth assistance. The experiments were performed at MOI=1 (i) and 0.1 (j), respectively. Data came from 3 independent experiments. The % of dependence was compared between the Ctrl and intrOv groups using ANOVA and Wilcoxon, respectively.

## Discussion

IntrOv is attenuated in genital tract pathogenicity (22, 23). However, the mechanisms of its attenuation in genital pathogenicity remain largely unknown. Prior studies of intrOv genital tract infection used an inoculating dose of 2 x 10^5^ IFUs. Here, we re-evaluated intrOv’s infectivity in the genital tract using different inoculating doses and by monitoring the live organism yields at the early stages of the infection. These studies led us to conclude that the ability of intrOv to productively infect the genital tract is impaired. First, the peak organism recovery from vaginal swabs following inoculation with 2 x 10^5^ IFU was on day 7 with intrOv compared to day 3 with the control. Second, intrOv recovery on days 1 and 3 was significantly lower than the control. Third, at an inoculum dose of 2 x 10^3^ IFUs, intrOv was no longer able to establish a productive infection. Fourth, intrOv organism recovery from lower genital tissues was lower than the control on day 3 post-infection. Fifth, the ascension of intrOv from the LGT to the UGT was impaired compared to the control. Finally, the ability of intrOv to infect HeLa cells *in vitro* was more dependent on centrifugation compared to the control, suggesting that intrOv has an attachment or entry defect. Efforts are underway to investigate how chlamydial virulence factors mutated in intrOv may promote *C. muridarum* to efficiently establish a productive infection in the lower genital tract.

The current study has revealed that wild-type control *C. muridarum* organisms can develop typical dose-dependent temporospatial infection courses in the female genital tract when the inoculation dose is within the range of 2 x 10^5^ to 2 x 10^3^ IFUs. A higher inoculation dose allowed *C. muridarum* to reach its peak infection earlier. The level of peaking infection also increased as the inoculation dose increased. Nevertheless, the length of the infection course remained similar between mice with different inoculation doses, suggesting that within this inoculation dose range, sufficient chlamydial infectivity was maintained without dramatically altering host responses. Extremely high or low inoculating doses significantly shortened chlamydial infection time courses in the female mouse genital tract due to the dramatically altered host responses or the lack of sufficient infection (27–29). Thus, the inoculation dose range used in the current study is suitable for investigating the biology of chlamydial interactions with the female genital mucosal tissues. Using this dose range, we compared the infectivity of the mutant intrOv with its isogenic wild-type *C. muridarum*, which led to the finding that intrOv is significantly deficient in infecting the female genital tract. The intrOv’s infection peaks were delayed considerably when the inoculating dose was high enough for intrOv to establish a productive infection or intrOv failed to establish an infection when the inoculating dose was reduced. This striking phenotype was not observed in previous studies in which the inoculation was used at or higher than 2 x 10^5^ IFUs (22, 23).

The significantly reduced titers of intrOv were mainly restricted to days 1 and 3, and by day 7, the intrOv’s titers reached the level of its control, indicating that intrOv’s deficiency in infectivity is transient. The delayed infectivity phenotype of intrOv is consistent with the observations that intrOv is less efficient in invading epithelial cells but maintains the same level of intracellular growth as its wild-type control once it enters an epithelial cell. A comparison of live chlamydial organisms recovered from different genital tissue segments revealed that the intrOv’s deficiency in infectivity was mainly restricted to the lower genital tract, suggesting that intrOv is either more susceptible to the host inhibition or less efficient in invading epithelial cells in the lower genital tract or both. More experiments are required to distinguish these possibilities. The current study has provided the experimental conditions for guiding the design of the new experiments and further revealing the mechanisms by which the mutated gene products in intrOv promote *C. muridarum* infectivity in the lower genital tract.

Previous studies have shown that intrOv is inhibited and cleared by IFNγ^+^ILC3s in the colon where its wild-type control maintains long-lasting colonization (30–32). It was found that both the wild-type control and the mutant intrOv *C. muridarum* organisms induced the intestinal IFNγ^+^ILC3s responses while only the wild-type but not intrOv could evade the IFNγ^+^ILC3s-mediated immunity. The question is whether the intrOv’s delay in establishing a productive infection in the lower genital tract tissues is also caused by IFNγ^+^ILC3s. A previous study reported that *C. trachomatis* induced IFNγ^+^ILC3s in the genital tract for inhibiting its own infection in the endometrial tissue (33). We hypothesize that intrOv may be able to induce IFNγ^+^ILC3s in the lower genital tract to delay the infection process. The chlamydial induction of colonic IFNγ^+^ILC3s depended on the IL-23 receptor signaling (32). Consistently, *C. muridarum* is known to induce IL-23 in the mouse genital tract (34, 35). Thus, we hypothesize that the intrOv-induced IL-23-IFNγ^+^ILC3s pathway may be responsible for delaying the intrOv infectivity in the lower genital tract. If so, it will be worth addressing why the intrOv-induced IL-23- IFNγ^+^ILC3s pathway completely blocks the infectivity of intrOv in the colon while only delaying/slowing the intrOv infection in the genital tract.

Finally, does the transient delay in intrOv’s establishing a productive infection in the lower genital tract contribute to its attenuation in the upper genital tract pathogenicity? IntrOv is highly attenuated in inducing hydrosalpinx following an intravaginal inoculation at an inoculating dose of 2 x 10^5^ IFUs (22). Although the intrOv’s attenuation in pathogenicity correlated with its inability to colonize the gastrointestinal tract (36), we now hypothesize that the slowed/delayed/reduced infectivity of intrOv in the LGT tissues may present a qualitatively different environmental cue or an attenuated cue to the mouse’s immune system, leading to an immune imprint that is less pathogenic compared to the more pathogenic immune imprints induced by the wild-type control organisms. Although intrOv eventually caught up with the control in infectious loads by day 7, the immune imprints developed before day 7 may remain active throughout the infection courses to regulate chlamydial pathogenicity in the upper genital tract. For example, chlamydial antigen-specific CD8^+^ T cells primed during the 1^st^ week of the control organism infection may promote chlamydial pathogenicity in the upper genital tract (11–13), while non-specific CD8^+^ T cells activated by intrOv may reduce chlamydial induction of hydrosalpinx (14, 15). More experiments are required to compare the early immune response profiles induced by intravaginal infection with intrOv or its control and to determine whether the intrOv-induced immune imprints can prevent chlamydial pathogenicity, while the wild-type control-induced imprints may exacerbate it.

## Materials and Methods

### 1. *Chlamydia* organisms

The *Chlamydia muridarum*mutant clone G28.51.1 (Genome seq: https://www.ebi.ac.uk/biosamples/samples/SAMN03273902) and its isogenic wild-type *C. muridarum* clone G13.32.1 (as a control) were used in the current study (22, 23). Due to its lack of pathogenicity in the genital tract (22, 23), the mutant clone G28.51.1 has been proposed as an *intr*acellular *O*ral vaccine *v*ector or intrOv (5). IntrOv contains two mutations: a substitutional mutation in gene *tc0237* leading to the 117^th^ codon change from glutamine (Q) to glutamic acid (E) or TC0237Q117E and a deletion mutation in *tc0668,* resulting in a stop codon at 216 that originally codes for glycine (G) or TC0668G216*. The intrOv and its wild-type control *C. muridarum* organisms were grown in HeLa cells (human cervical carcinoma epithelial cells; ATCC# CCL-2), and density gradient centrifugation was used to purify elementary bodies (EBs). The purified EBs were stored in aliquots @ −80°C until use.

### 2. Mouse infection

The mouse experiments were carried out following the recommendations in the Guide for the Care and Use of Laboratory Animals endorsed by the National Institutes of Health. The protocol was approved by the Committee on the Ethics of Laboratory Animal Experiments of the University of Texas Health Science Center at San Antonio.

Five to 7 week-old female mice (C57BL/6J, stock No: 000664) were purchased from Jackson Laboratories, Inc., Bar Harbor, ME. All mice were intravaginally infected with intrOv or its wild-type control EBs at the inoculating doses of 2 × 10^5^, 2 × 10^4^, or 2 × 10^3^ inclusion forming units (IFUs) per mouse, as indicated in individual experiments. Briefly, EBs diluted in 10 μl SPG (220 mM sucrose, 12.5 mM phosphate, 4 mM l-glutamic acid, pH 7.5) buffer were delivered to the upper vaginal cavity of each mouse using a p20 micropipette. The mouse was held upside down with the vagina facing the operator during the inoculation. Efforts were made to deliver the 10 μl to the space between the ectocervix and one side of the vaginal wall. After ejecting the inoculum, slowly pull out the inoculation tip and hold the mouse in the inoculation position for 3 to 5 minutes to reduce potential leaking. To increase the mouse susceptibility to chlamydial infection and ensure inoculation reproducibility, mice were primed by intraperitoneal injection with 2.5 mg of colloidal depot medroxyprogesterone (Depo-Provera; Pharmacia & Upjohn LLC, Kalamazoo, MI) suspended in sterile phosphate-buffered saline (PBS) 7 days before the intravaginal infection. In addition, all mice were pre-swabbed to remove any vaginal plugs before the intravaginal inoculation. After the inoculation, mice were monitored for live organism shedding in vaginal swabs or sacrificed for titrating live organisms in the genital tract tissues.

### 3. Titrating live chlamydial organisms from mouse swabs and tissues

To monitor live chlamydial organisms from the genital tracts, vaginal swabs were collected in 0.5 ml of SPG buffer and vortexed with glass beads to release infectious EBs. For titrating live organisms from mouse genital tract tissues, the lower genital tract (LGT) tissues, including the lower vagina (LV), upper vagina (UV), and ectocervix (EC), and the upper genital tract (UGT) tissues, including endocervix (NC), uterine horns from both sides (UH), and Oviducts/ovaries (OV) from both sides, were collected from mice sacrificed on different days after the intravaginal inoculation as indicated in individual experiments. Each tissue segment was collected to 0.5 ml SPG buffer, followed by homogenization and sonication. Live organisms in the supernatants were titrated on HeLa cells in duplicate. The total number of IFUs per swab or tissue was converted into log_10_ for calculating the group mean and standard deviation. Please note that the detection limits of the titration method are 10 IFUs per swab and 40 IFUs per tissue sample. A total of 100 μl from each sample without (neat) or with serial dilutions was used to inoculate monolayer HeLa cells. After incubation, the inoculated culture wells were processed for immunofluorescence labeling, and the chlamydial inclusions were counted to calculate the total number of live organisms recovered from each sample.

### 4. Immunofluorescence assay

The immunofluorescence assay for visualizing and counting chlamydial inclusions in Chlamydia-infected HeLa cultures was described previously (37). Briefly, infected HeLa cells grown on 96-well plates were fixed with paraformaldehyde (Sigma, St. Louis, MO 63178) and permeabilized with 0.1% Triton X-100 (Thermo Scientific, A16046.0F). The monolayers were labeled with a rabbit anti-chlamydial antibody (raised by immunization with *C. muridarum* EBs) and a goat anti-rabbit IgG conjugated with fluorescein isothiocyanate (FITC, green, Jackson ImmunoResearch Laboratories, Inc) to visualize chlamydial inclusions while a Hoechst dye (blue; Sigma) was used for labeling nuclear DNA. The labeled cells were viewed under an Olympus IX-80 fluorescence microscope equipped with multiple filter sets (Olympus, Melville, NY).

### 5. DEAE/Centrifugation-assisted attachment/entry dependence assay

Cell culture infectivity of *C. muridarum* organisms was measured by comparing their dependence on the treatment of HeLa cells with DEAE (diethylaminoethyl dextran, cat# D9885-50G, Sigma-Aldrich, Inc. St. Louis, MO) and centrifuging the inoculum onto HeLa monolayers during *in vitro* infection. Purified *C. muridarum* EBs were serially diluted in SPG before being inoculated onto confluent HeLa monolayers grown in 96- well tissue culture plates in duplicate. The previously described assisted attachment and entry infection conditions (6, 7) were used, including DEAE-dextran (in DMEM medium without serum) pretreatment of HeLa monolayers for 10 min at 37°C before inoculation. After removing the DEAE solution, serially diluted inoculum in 100 μl SPG was inoculated to each well, followed by centrifugation at 1000 rpm (214.7g, Sorvall 75006445 rotor, Inc. Thermo Fisher Scientific, Waltham, MA) for 1h at room temperature. To remove either assisting condition, the DEAE solution was replaced with the diluting DMEM medium alone (-DEAE), the inoculum-containing culture plates were kept at the same temperature without centrifugation (-Centrifuge), or both (-DEAE & - Centrifuge). After completing the attachment/entry steps, the culture plates were incubated in DMEM medium containing 2 μg/ml of cycloheximide (Cat# C7698-5G, Sigma-Aldrich, Inc. St. Louis, MO) at 37°C for 24h before being fixed and processed for immunofluorescence detection of *C. muridarum* inclusions.

### 6. Cycloheximide-assisted intracellular growth dependence assay

Chlamydial dependence on the intracellular growth assistance provided by cycloheximide was measured by comparing the IFUs detected from cultures grown in DMEM medium with or without cycloheximide following the same infection procedures described above.

The results from the above attachment/entry/intracellular growth assistance experiments were expressed as % of dependence on a given assistance condition for growth. The % of dependence was calculated in two steps, including normalization and % of dependence: To normalize, the IFUs from the complete assistance culture were used as the denominator to divide the IFUs from the culture that is absent of an assistance condition. The normalized result is the % of remaining growth in the absence of a given assistance. To obtain the % of dependence on a given assisting condition, the corresponding % of remaining growth was subtracted from 100%.

### 7. Statistics

The numbers of live organisms in IFUs at individual data points or over a time course were compared using the Wilcoxon rank-sum test. Area-under-the-curve or AUC was used for comparing the time course or clusters of tissue sample data. When multiple groups were included in a given experiment, ANOVA was first used to determine whether there was an overall significant difference among all groups. Only when p<0.05 (ANOVA) were the differences between each two groups further analyzed using Wilcoxon.

